## Supplementary Information for "Hexokinase-I directly binds to a charged membrane-buried glutamate of mitochondrial VDAC1 and VDAC2"

**This PDF file includes:**

Supplementary Figs. S1 to S6

Movies S1 and S2

Supplementary Tables S1 to S3

Image J Macro

Unprocessed images of immunoblots

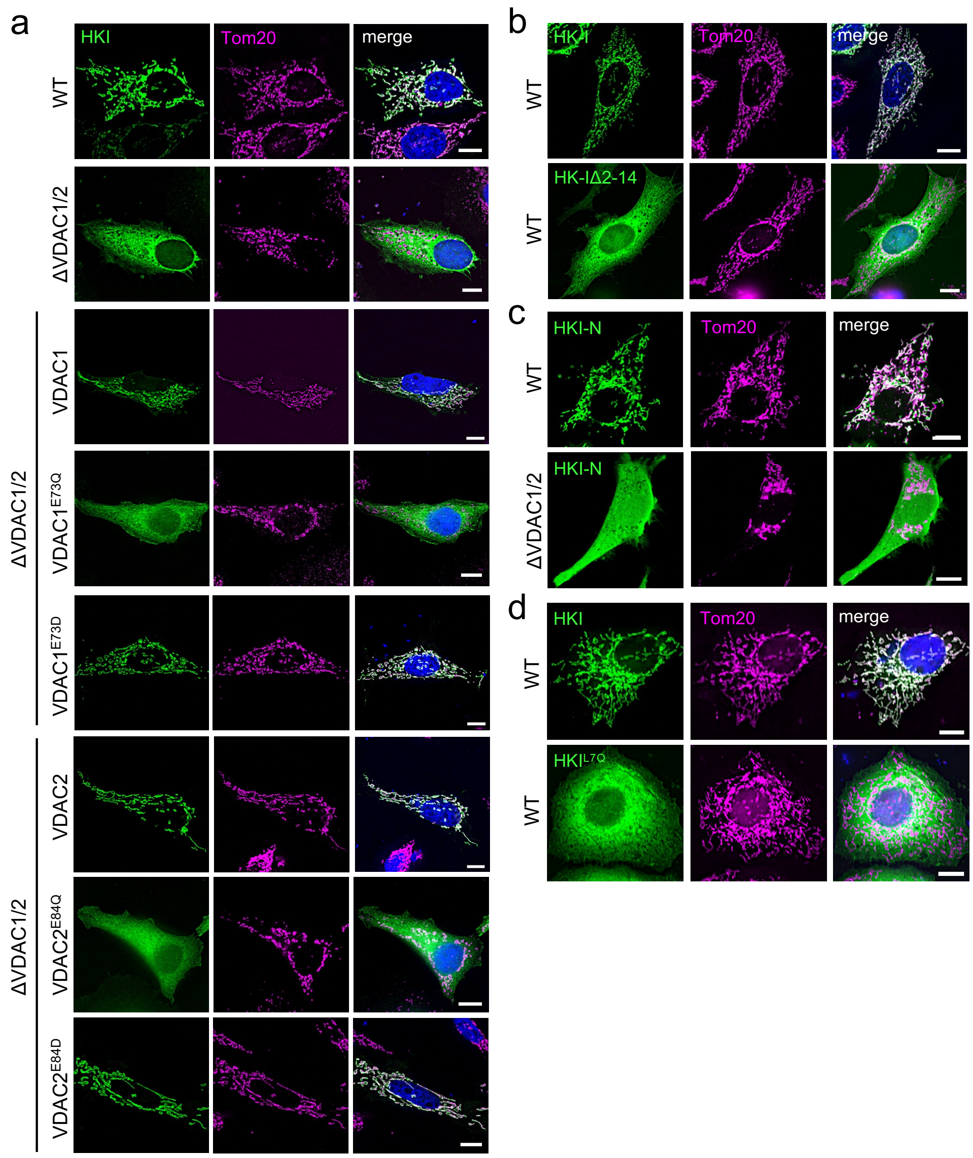


**Figure S1 | Mitochondrial localization of HKI relies on its N-terminal α-helix and a membrane-buried Glu in VDACs.** (**a**) Fluorescence images of wild-type (WT) and VDAC1/2-DKO HeLa cells expressing EGFP-tagged HKI (*green*) alone or in combination with HA-tagged VDAC1, VDAC1$^{E73Q}$, VDAC1$^{E73D}$, VDAC2, VDAC2$^{E84Q}$ or VDAC2$^{E84D}$, fixed and then stained with DAPI (*blue*) and an antibody against Tom20 (*magenta*). Scale bar, 10 μm. (**b**) Fluorescence images of WT HeLa cells expressing EGFP-tagged HKI or *N*-terminal truncation mutant HKI$_{\Delta 2\text{-}14}$, fixed and then stained with DAPI (*blue*) and an antibody against Tom20 (*magenta*). Scale bar, 10 μm. (**c**) Fluorescence images of live WT and VDAC1/2-DKO HeLa cells co-expressing EGFP-tagged Tom20 (*magenta*) and Halo-tagged HKI-N (*N*-terminal HKI residues 1-17, *green*). Scale bar, 10 μm. (**d**) Fluorescence images of WT HeLa cells expressing EGFP-tagged HKI or HKI$^{L7Q}$ (*green*), fixed and then stained with DAPI (*blue*) and an antibody against Tom20 (*magenta*). Scale bar, 10 μm.

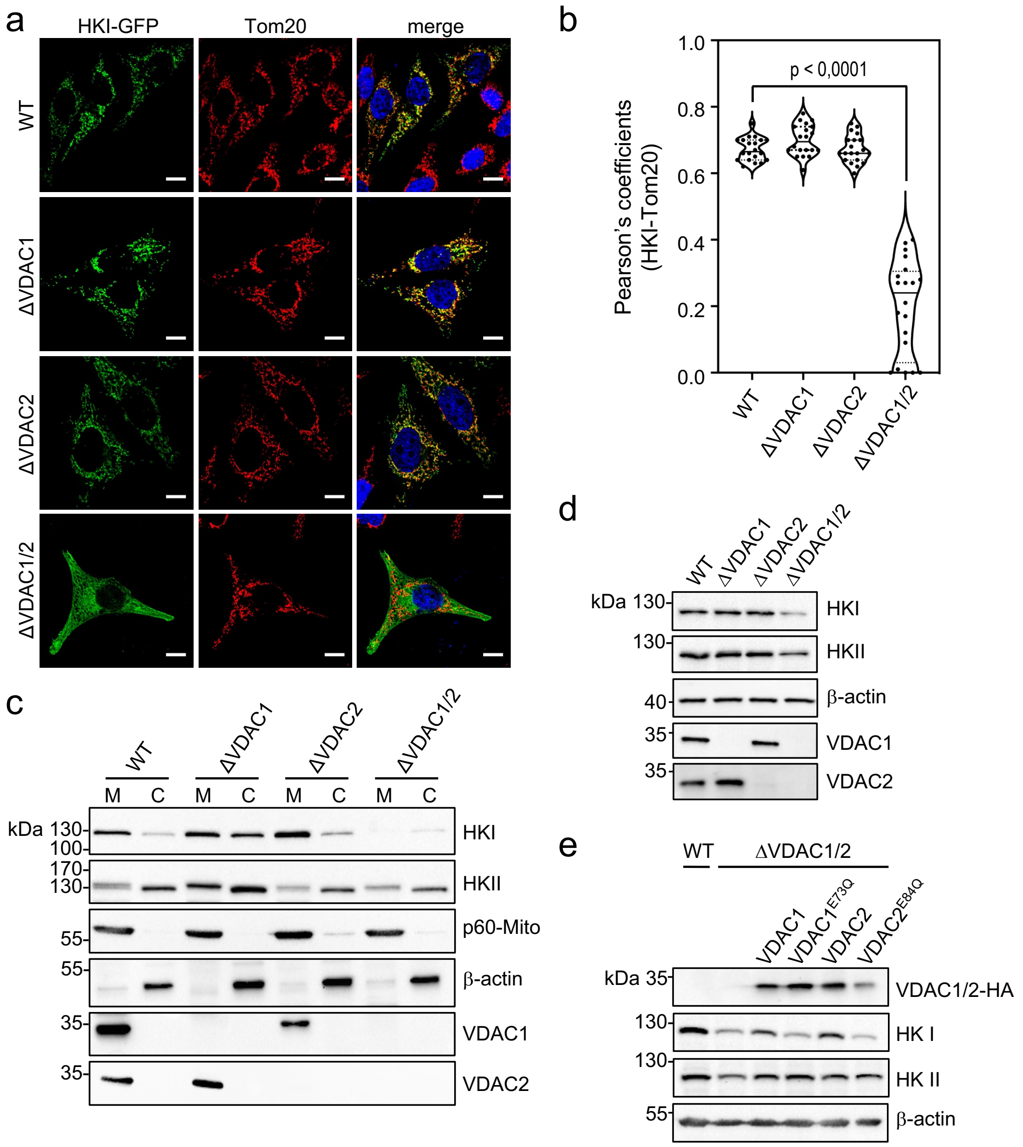


**Figure S2 | Both VDAC1 and VDAC2 contribute to stabilizing the mitochondrial pool of HKI.** (**a**) Fluorescence images of wild-type (WT) and VDAC1-KO, VDAC2-KO and VDAC1/2-DKO HeLa cells expressing EGFP-tagged HKI (*green*) fixed and then stained with DAPI (*blue*) and an antibody against Tom20 (*red*). Scale bar, 10 μm. (**b**) Pearson's correlation co-efficient analysis between HKI and Tom20 signals in cells as in (a). Data shown are based on the analysis of 20 cells per condition from two independent experiments. *P* values were calculated by unpaired two-tailed *t* test. (**c**) WT, VDAC1-KO, VDAC2-KO and VDAC1/2-DKO HeLa cells were subjected to subcellular fractionation. Mitochondrial (M) and cytosolic fractions (C) were analysed by immunoblotting with antibodies against VDAC1, VDAC2, HKI, HKII, p60-Mito and β-actin. (**d**) Total cell lysates from WT, VDAC1-KO, VDAC2-KO and VDAC1/2-DKO human HCT116 cells were subjected to immunoblotting with antibodies against VDAC1, VDAC2, HKI, HKII and β-actin. (**e**) Total cell lysates from WT and VDAC1/2-DKO human HCT116 cells transduced with HA-tagged VDAC1, VDAC1$^{E73Q}$, VDAC2 or VDAC2$^{E84Q}$ were subjected to immunoblotting with antibodies against the HA-tag, HKI, HKII and β-actin. Immunoblots stained with anti-β-actin, anti-VDAC1 and anti-VDAC2 antibodies in (d) and (e) are identical to those shown in Suppl. Fig. S10 of Dadsena et al.[1]

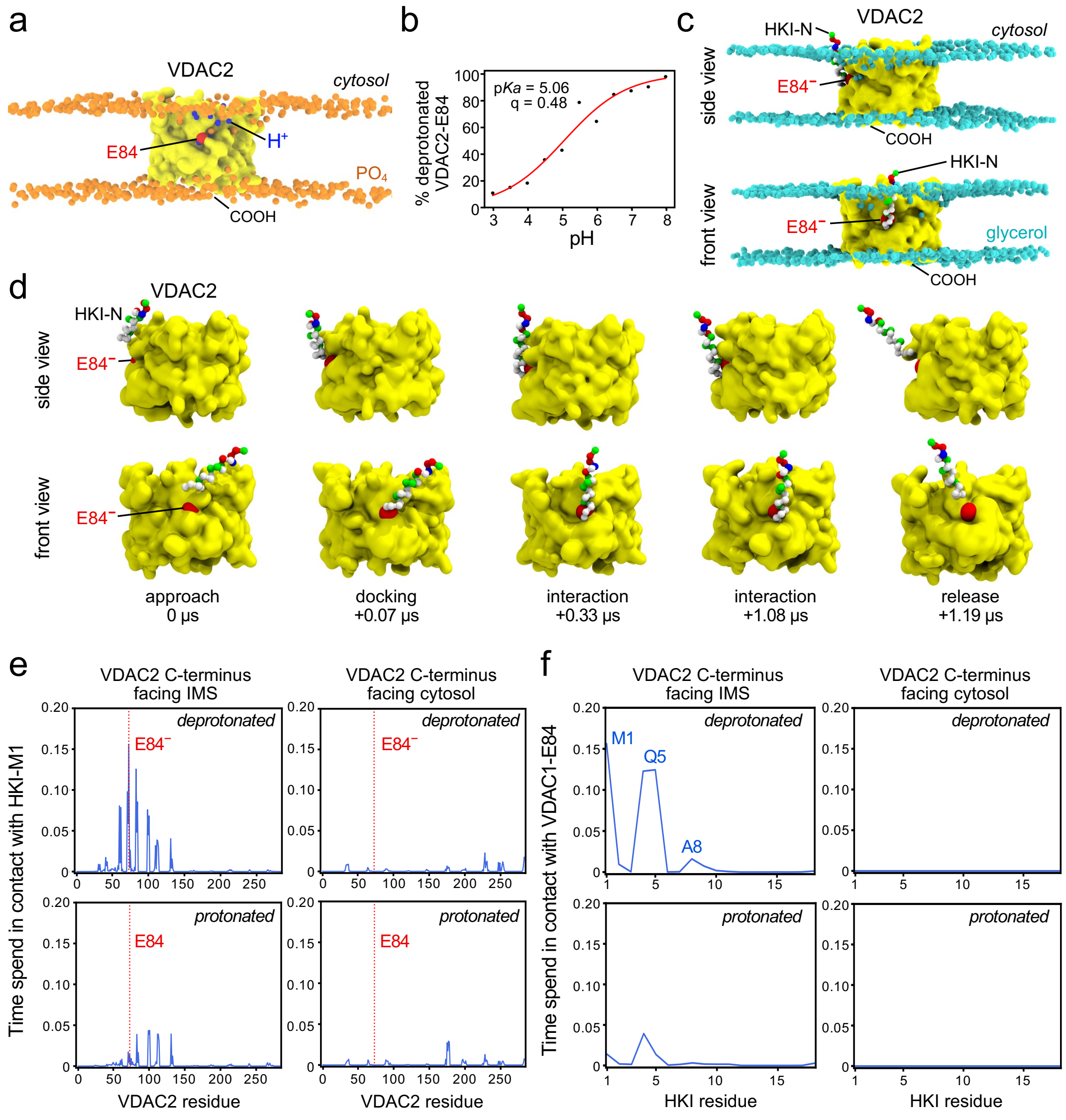


**Figure S3 | HKI-N binding to VDAC2 is directly controlled by the protonation state of the membrane-buried Glu.** (**a**) Still from a titratable MD simulation of VDAC2 (*yellow*) to evaluate the protonation state of E84 (*red*) at pH 5.0. $PO_4$ groups in the POPC-based bilayer are marked in *orange* and protons are marked in *blue*. (**b**) Titration curve showing the degree of deprotonation of E84 in VDAC2, simulated at a pH range of 3-8. (**c**) Stills from an MD simulation showing HKI-N bound to VDAC2 with a deprotonated E84 (*red*) and IMS-facing *C*-terminus. Glycerol groups in the OMM-mimicking bilayer are marked in *cyan*. (**d**) Stills from an MD simulation, showing the approach and binding of HKI-N to VDAC2 with a deprotonated E84 (*red*) and IMS-facing *C*-terminus. (**e**) Relative duration of contacts between HKI-Met1 and specific residues of VDAC2 with a protonated or deprotonated E84 and cytosol- or IMS-facing *C*-terminus. Shown are the combined data of three individual replicas with a total simulation time between 168 µs and 177 µs per condition. (**f**) Relative duration of contacts between VDAC2-E84 and specific residues of HKI-N under the same conditions as in (e).

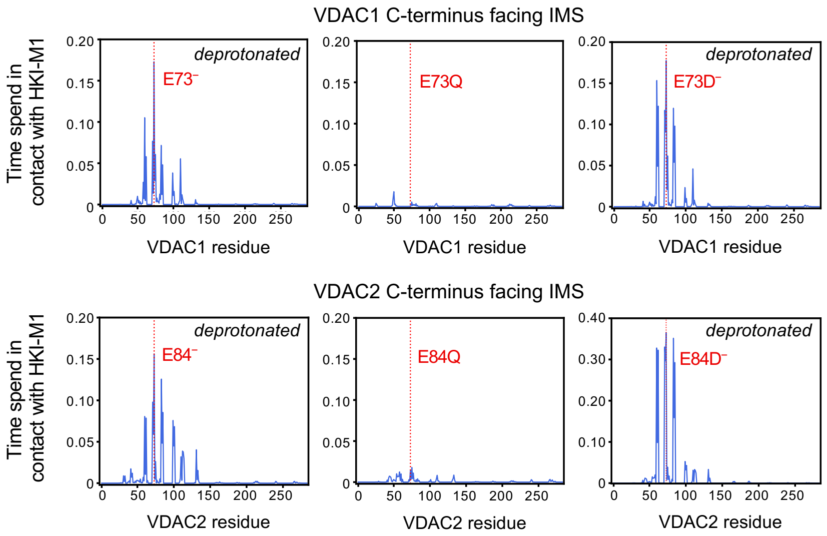


**Figure S4 | VDACs with a Glu-to-Asp substitution retain the ability to bind HKI-N.** Relative duration of contacts between HKI-Met1 and specific residues of VDAC1, VDAC1$^{E73Q}$, VDAC1$^{E73D}$, VDAC2, VDAC2$^{E84Q}$ and VDAC2$^{E84D}$ with deprotonated bilayer-facing acidic residues and IMS-facing *C*-termini simulated in an OMM-mimicking bilayer. Shown are the combined data of three individual replicas with a total simulation time of 163 μs and 211 μs per condition.

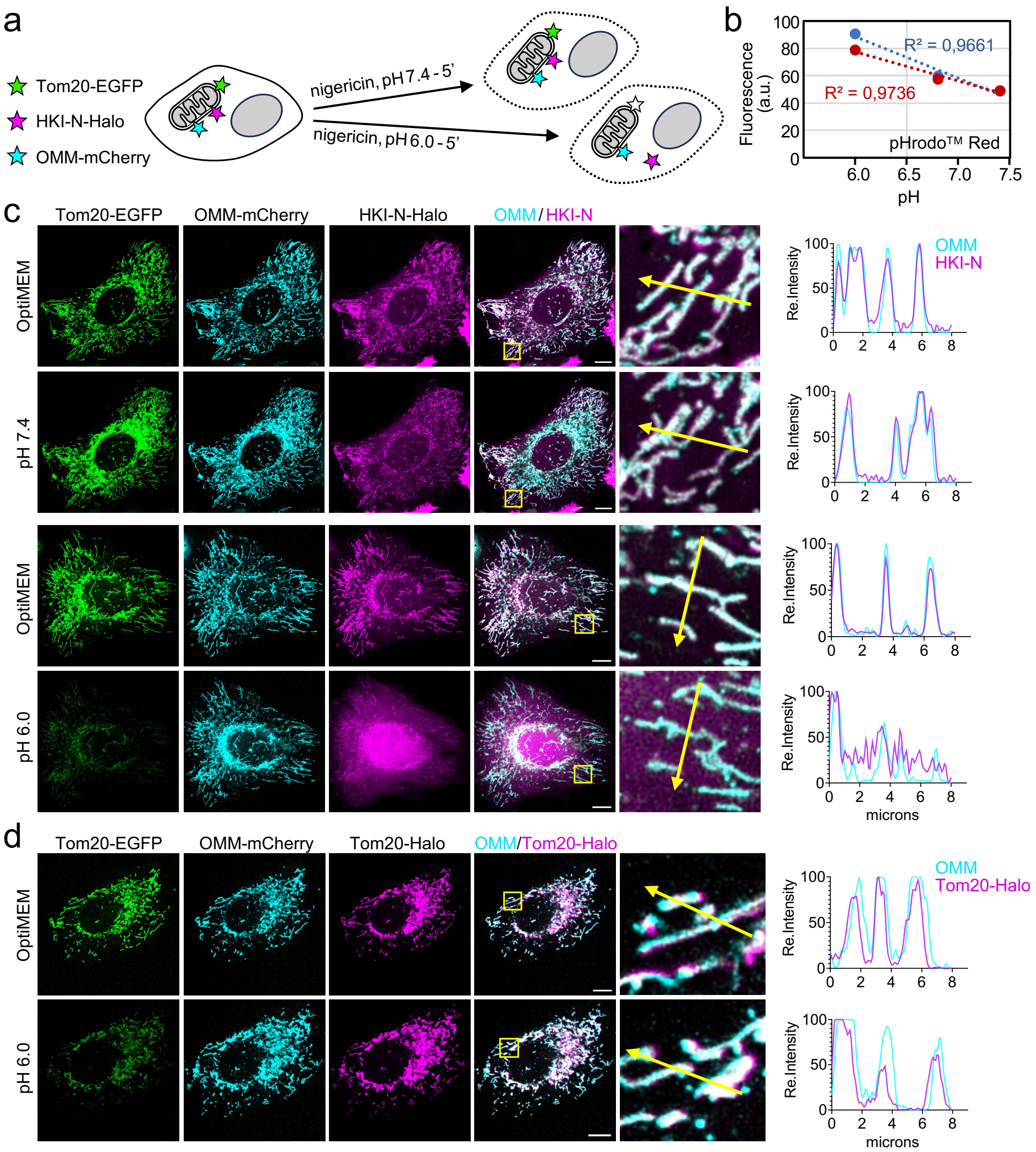


**Figure S5 | Acidification of cytosolic pH triggers dissociation of HKI-N from mitochondria.** (**a**) Schematic outline of experimental strategy to determine the impact of cytosolic acidification on mitochondrial association of HKI-N. (**b**) HeLa cells were incubated with 5 µM pHrodo™ Red AM and 10 µM nigericine in buffers with descending pH (7.4, 6.8 and 6.0) for 10 min at 37°C per buffer. Relative fluorescence levels were plotted in the graph and a linear trendline was fitted to get the pH standard curve. Data shown are from two independent experiments. (**c**) Fluorescence images of live HeLa cells co-expressing EGFP-tagged Tom20 (*green*), OMM-mCherry (*cyan*) and Halo-tagged HKI-N (*magenta*) grown in Optimem (top) and then treated with nigericin in pH 7.4 buffer or pH 6.0 buffer for 5 min. Line scans showing degree of overlap between OMM and HKI-N signals along the path of the arrow shown in the zoom-in. Scale bar, 10 µm. (**d**) Fluorescence images of live HeLa cells co-expressing EGFP-tagged Tom20 (*green*), OMM-mCherry (*cyan*) and Halo-tagged Tom20 (*magenta*) grown in Optimem (top) and then treated with nigericin in pH 6.0 buffer for 5 min. Line scans showing degree of overlap between OMM and Tom20-Halo signals along the path of the arrow shown in the zoom-in. Scale bar, 10 µm.

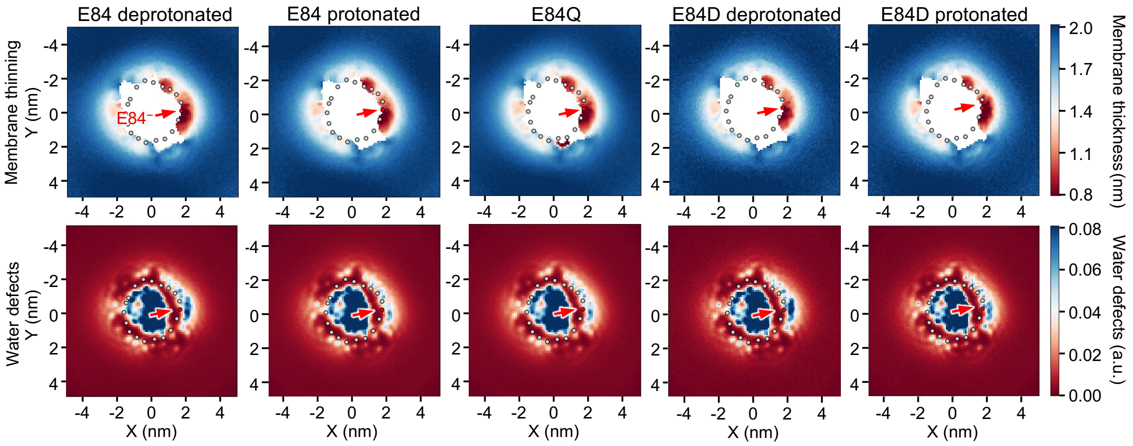


**Figure S6 | VDAC2 causes lipid packing defects and membrane leaflet thinning proximal to the bilayer-facing Glu.** Cytosolic leaflet thinning and water defect graphs of VDAC2, VDAC2$^{E84Q}$ and VDAC2$^{E84D}$ simulated in a DOPC bilayer with *C*-termini facing the IMS (bottom) leaflet. The bilayer-facing acidic residues were protonated or deprotonated, as indicated. Analysis was done as in Fig. 6a and 6b.

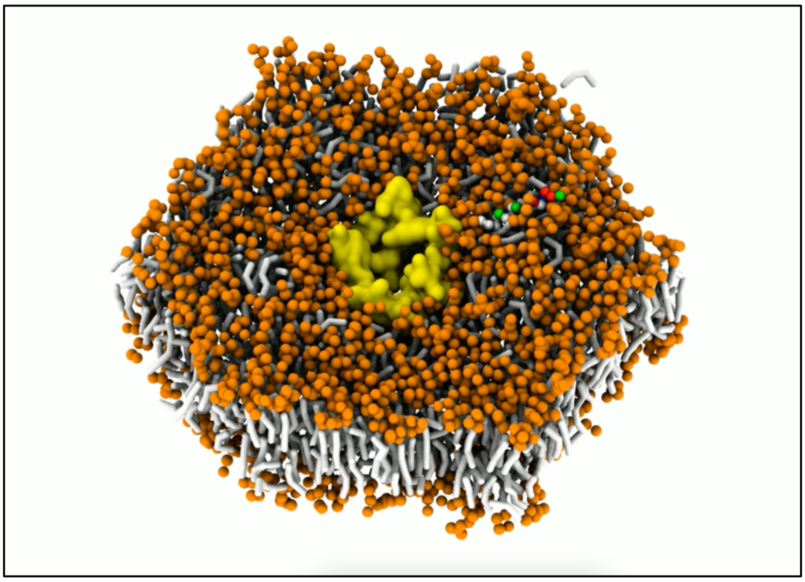

**Supplementary Movie 1 | MD simulation of complex formation between HKI-N and VDAC1.**
Simulation of HKI-N binding to VDAC1 with a deprotonated E73 and IMS-facing *C*-terminus in an OMM-mimicking bilayer. The movie captures 849 ns of simulated time. The image is a screen shot of the first frame of the movie.

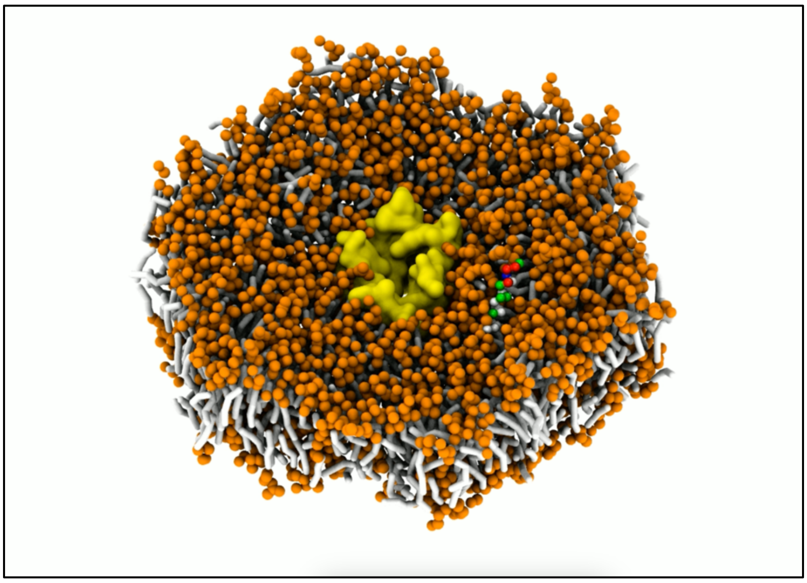

**Supplementary Movie 2 | MD simulation of complex formation between HKI-N and VDAC2.**
Simulation of HKI-N binding to VDAC2 with a deprotonated E84 and IMS-facing *C*-terminus in an OMM-mimicking bilayer. The movie captures 1400 ns of simulated time. The image is a screen shot of the first frame of the movie.

**Table S1 | Simulated times per system**

| System | Box size | Simulation time |
|---|---|---|
| **VDACs with HKI-N** | | |
| VDAC1 E73 deprotonated $COOH_{IMS}$ | x=16nm, y=14nm, z=9nm | 210.9µs |
| VDAC1 E73 protonated $COOH_{IMS}$ | x=16nm, y=14nm, z=9nm | 169.3µs |
| VDAC1 E73 deprotonated $COOH_{cyto}$ | x=16nm, y=14nm, z=9nm | 169.4µs |
| VDAC1 E73 protonated $COOH_{cyto}$ | x=16nm, y=14nm, z=9nm | 177.9µs |
| VDAC2 E84 deprotonated $COOH_{IMS}$ | x=16nm, y=14nm, z=9nm | 170.3µs |
| VDAC2 E84 protonated $COOH_{IMS}$ | x=16nm, y=14nm, z=9nm | 170.4µs |
| VDAC2 E84 deprotonated $COOH_{cyto}$ | x=16nm, y=14nm, z=9nm | 168.9µs |
| VDAC2 E84 protonated $COOH_{cyto}$ | x=16nm, y=14nm, z=9nm | 177.1µs |
| **VDACs w/o HKI-N** | | |
| VDAC1 E73 deprotonated $COOH_{IMS}$ | x=11nm, y=11nm, z=15nm | 32.4µs |
| VDAC1 E73 protonated $COOH_{IMS}$ | x=11nm, y=11nm, z=15nm | 31.9µs |
| VDAC2 E84 deprotonated $COOH_{IMS}$ | x=11nm, y=11nm, z=15nm | 32.6µs |
| VDAC2 E84 protonated $COOH_{IMS}$ | x=11nm, y=11nm, z=15nm | 32.4µs |
| **VDAC mutants with HKI-N** | | |
| VDAC1 E73Q $COOH_{IMS}$ | x=16nm, y=14nm, z=9nm | 163.5µs |
| VDAC1 E73D deprotonated $COOH_{IMS}$ | x=16nm, y=14nm, z=9nm | 300.4µs |
| VDAC1 E73D protonated $COOH_{IMS}$ | x=16nm, y=14nm, z=9nm | 158.0µs |
| VDAC2 E84Q $COOH_{IMS}$ | x=16nm, y=14nm, z=9nm | 173.4µs |
| VDAC2 E84D deprotonated $COOH_{IMS}$ | x=16nm, y=14nm, z=9nm | 199.1µs |
| VDAC1 E73F/F71E $COOH_{IMS}$ | x=16nm, y=14nm, z=9nm | 162.8µs |
| VDAC1 E73F/F71E $COOH_{cyto}$ | x=16nm, y=14nm, z=9nm | 148.9µs |
| VDAC1 S101L $COOH_{IMS}$ | x=16nm, y=14nm, z=9nm | 172.4µs |
| VDAC1 T77L $COOH_{IMS}$ | x=16nm, y=14nm, z=9nm | 171.0µs |
| VDAC1 S101L/T77L $COOH_{IMS}$ | x=16nm, y=14nm, z=9nm | 169.7µs |
| **VDAC mutants w/o HKI-N** | | |
| VDAC1 E73Q $COOH_{IMS}$ | x=11nm, y=11nm, z=15nm | 55.2µs |
| VDAC1 E73D deprotonated $COOH_{IMS}$ | x=11nm, y=11nm, z=15nm | 49.6µs |
| VDAC1 E73D protonated $COOH_{IMS}$ | x=11nm, y=11nm, z=15nm | 30.1 µs |
| VDAC2 E84Q $COOH_{IMS}$ | x=11nm, y=11nm, z=15nm | 59.1µs |
| VDAC2 E84D $COOH_{IMS}$ | x=11nm, y=11nm, z=15nm | 57.2µs |
| VDAC1 E73F/F71E $COOH_{IMS}$ | x=11nm, y=11nm, z=15nm | 26.4µs |
| VDAC1 S101L $COOH_{IMS}$ | x=11nm, y=11nm, z=15nm | 17.9µs |
| VDAC1 T77L $COOH_{IMS}$ | x=11nm, y=11nm, z=15nm | 27.7µs |
| VDAC1 S101L/T77L $COOH_{IMS}$ | x=11nm, y=11nm, z=15nm | 27.4µs |
| **Total simulation time: 3,713.3µs** | | |

**Table S2 | Primers used in this study**

| Primer name | Primer sequence (5'-3') |
|---|---|
| pSEMS HKI-N-Halo fwd | GCTGAAGATGATGTG GAAGTGTGGTGGAATTCCATGATCG |
| pSEMS HKI-N-Halo rev | CCTGTGCCATGGCCCCCGTCATCCTTCAGCTCCG |
| pEGFP ratHKI-L7Q fwd | CGCGCAACTACAGGCCTATTACT |
| pEGFP ratHKI-L7Q rev | GCGATCATGCTGACGGTG |
| pEGFP ratHKIΔ2-14 fwd | AAGGATGACCAAGTCAAAAAGATTGACAAG |
| pEGFP ratHKIΔ2-14 rev | CATGCTGACGGTGGGGGA |
| pcDNA3.1 hVDAC1-HA fwd | AAGCTGGCTAGCACCATGGCAATGGCTGTGCCACCCACG |
| pcDNA3.1 hVDAC1-HA rev | GGGCCCTCTAGATCAGGCGTAATCCGGCACATCATAGGGG TATGCTTGAAATTCCAGTCC |
| pcDNA3.1 hVDAC2-HA fwd | AAGCTGGCTAGCACCATGGCAATGGCGACCCACGGACAG |
| pcDNA3.1 hVDAC2-HA rev | GGGCCCTCTAGATCAGGCGTAATCCGGCACATCATAGGGG TAAGCGTCCAACTCCAGGGC |
| pcDNA3.1 hVDAC1$^{E73Q}$-HA fwd | GGCCTGACGTTTACACAGAAATGG AATACCGAC |
| pcDNA3.1 hVDAC1$^{E73Q}$-HA rev | GTCGGTATTCCATTTCTGTGTAAACGTCAGGCC |
| pcDNA3.1 hVDAC2$^{E84Q}$-HA fwd | GGTCTGACTTTCACACAAAAGTGGAACACTGATAAC |
| pcDNA3.1 hVDAC2$^{E84Q}$-HA rev | GTTATCAGTGTTCCACTTTTGTGTGAAAGTCAGACC |
| pcDNA3.1 hVDAC1$^{E73D}$-HA fwd | GACAAATGGAATACCGACAATACACTAG |
| pcDNA3.1 hVDAC1$^{E73D}$-HA rev | TGTAAACGTCAGGCCGTACTCAG |
| pcDNA3.1 hVDAC2$^{E84D}$-HA fwd | GACAAGTGGAACACTGATAACACTCTG |
| pcDNA3.1 hVDAC2$^{E84D}$-HA rev | TGTGAAAGTCAGACCATACTCAC |
| pcDNA3.1 hVDAC1$^{E73F/F71E}$-HA fwd | GAGACATTTAAATGGAATACCGACAATACACTAG |
| pcDNA3.1 hVDAC1$^{E73F/F71E}$-HA rev | CGTCAGGCCGTACTCAGTCC |
| pcDNA3.1 hVDAC1$^{T77L}$-HA fwd | CTGGACAATACACTAGGCACCGAG |
| pcDNA3.1 hVDAC1$^{T77L}$-HA rev | ATTCCATTTCTCTGTAAACGTCAG |
| pcDNA3.1 hVDAC1$^{S101L}$-HA fwd | CTGTCCTTCTCACCTAACACTGGG |
| pcDNA3.1 hVDAC1$^{S101L}$-HA rev | ATCGAAGGTCAGCTTCAGTCC |
| pcDNA3.1 (+) VDAC1$^{T77L/S101L}$-HA fwd | CTGGACAATACACTAGGCACCGAG |
| pcDNA3.1 (+) VDAC1$^{T77L/S101L}$-HA rev | ATTCCATTTCTCTGTAAACGTCAG |

**Table S3 | Chromatic shifts in reference to GFP channel**

| Shift | mCherry | JF646 |
|---|---|---|
| Dx (µm) | -0.06 | -0.131 |
| Dy (µm) | -0.006 | 0.017 |
| Dz (µm) | -0.314 | -0.648 |
| Rot (°) | 0.034 | -0.038 |
| Scale | 0.999 | 1.001 |

**Image J Macro**

<u>Pearson's analysis</u>

```
dir1 = getDirectory("Choose Source Directory");
list = getFileList(dir1);
counter = 0;
var channel, slice, frame;

for (i=0; i<list.length; i++) {
filename = dir1 + list[i];
//check if the current file is in fact a subfolder
if (File.isDirectory(filename)) {
}
else {
if (endsWith(filename, ".dv")) {
run("Bio-Formats Importer", "open=["+filename+"] autoscale color_mode=Default rois_import=[ROI manager] view=Hyperstack stack_order=XYCZT");
selectImage(list[i]);
run("Maximize");
Stack.setPosition(3, 8, 0);
setTool("polygon");
waitForUser("Z and ROI", "Select a z-plane and draw ROI - polygon tool is already selected! Then click 'Ok'");
setBatchMode(true);
Stack.getPosition(channel, slice, frame);
run("Crop");
setBackgroundColor(0, 0, 0);
Stack.setChannel(3);
run("Duplicate...", "  title=channel2 channels=3 slices="+slice);
run("Clear Outside");
selectImage(list[i]);
Stack.setChannel(1);
run("Duplicate...", "  title=channel3 channels=1 slices="+slice);
run("Clear Outside");
close(list[i]);
run("JACoP ", "imga=channel2 imgb=channel3 pearson costesthr");
dataStr = split( getInfo("Log"), "\n" );
res = substring( dataStr[5], indexOf( dataStr[5], "=" ) + 1, lengthOf( dataStr[5] ) );
costes = substring( dataStr[9], indexOf( dataStr[9], "=" ) + 1, indexOf( dataStr[9], "(" ) - 1);
print("\\Clear");
```

```
setResult("Filename", counter, list[i]);
setResult("Pearson", counter, res);
setResult("Costes", counter, costes);
close("channel2");
close("channel3");
close('Costes*');
counter = counter+1;
setBatchMode(false);
}
}
}
selectWindow("Log");
run("Close");
```

### Supplementary reference


1. Dadsena, S. *et al.* Ceramides bind VDAC2 to trigger mitochondrial apoptosis. *Nat. Commun.* **10**, (2019).

### Unprocessed images of immunoblots

#### Figure S2c - uncropped

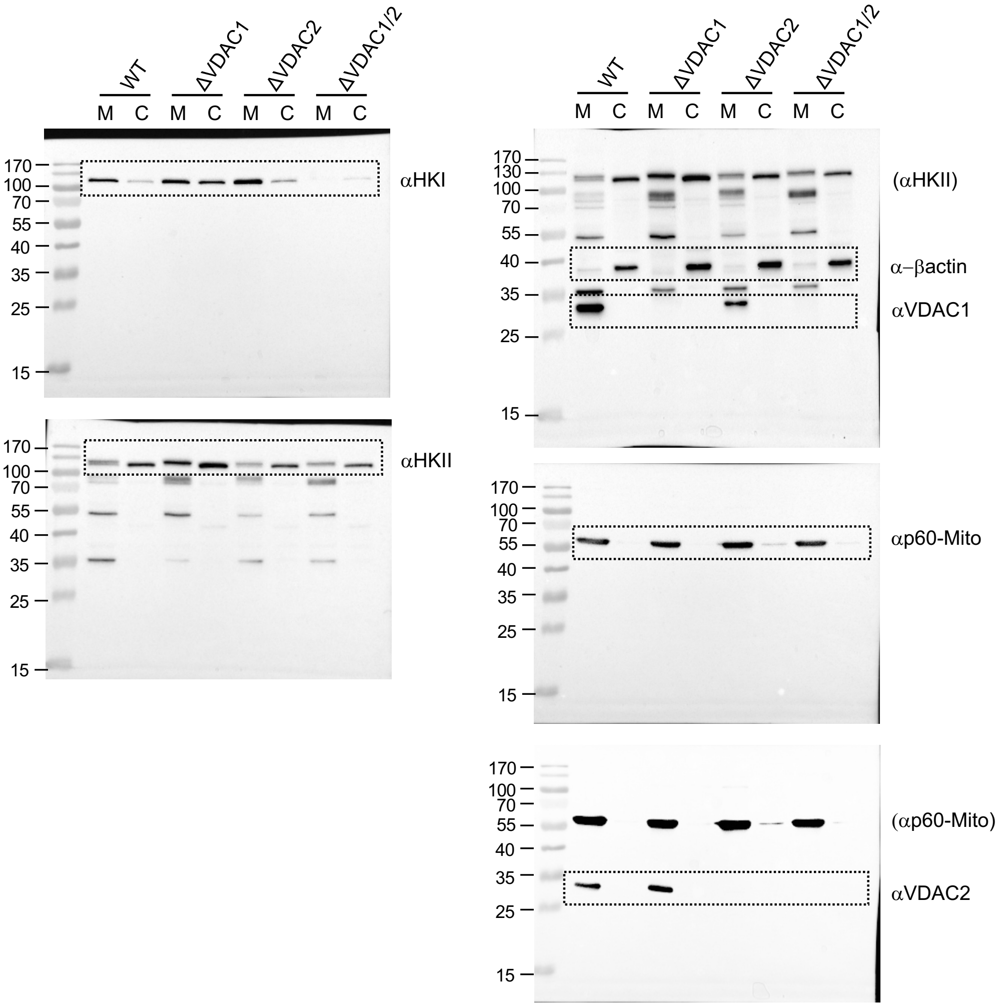

### Figure S2d - uncropped

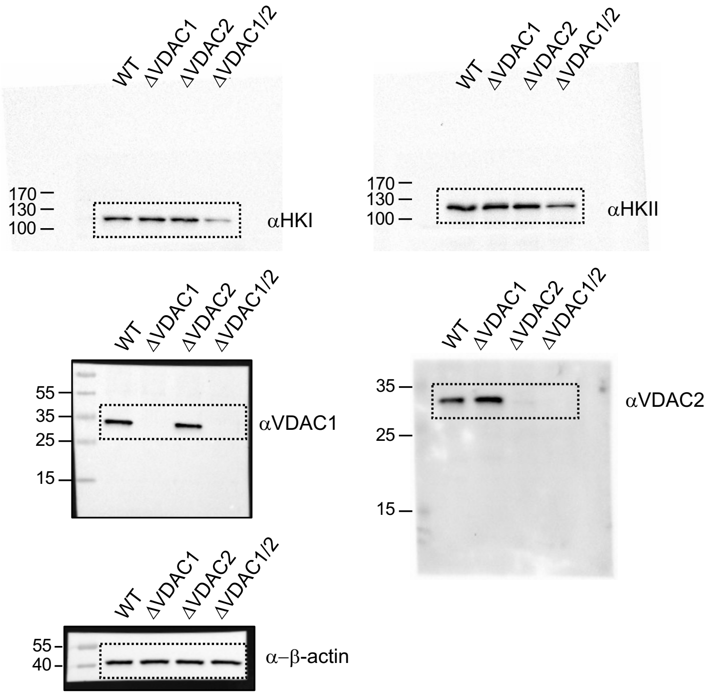


### Figure S2e - uncropped

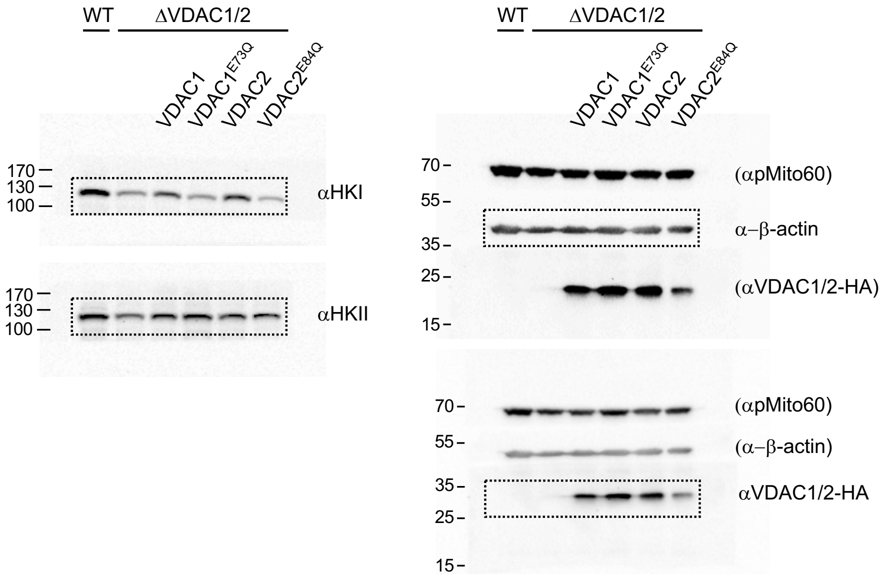
